# Does somatosensory acuity influence the extent of internal model recalibration in young and older adults?

**DOI:** 10.1101/2020.10.16.342295

**Authors:** Koenraad Vandevoorde, Jean-Jacques Orban de Xivry

## Abstract

The ability to adjust movements to changes in the environment declines with aging. This age-related decline is caused by the decline of explicit adjustments. However, automatic adjustment of movement, or internal model recalibration, remains intact and might even be increased with aging. Since somatosensory information appears to be required for internal model recalibration, it might well be that an age-related decline in somatosensory acuity is linked to the increase of internal model recalibration. One possible explanation for an increased internal model recalibration is that age-related somatosensory deficits could lead to altered sensory integration with an increased weighting of the visual sensory-prediction error. Another possibility is that reduced somatosensory acuity results in an increased reliance on predicted sensory feedback. Both these explanations led to our preregistered hypothesis: we expect a relation between the decline of somatosensation and the increased internal model recalibration with aging. However, we failed to support this hypothesis. Our results question the existence of reliability-based integration of visual and somatosensory signals during motor adaptation.

**New & Noteworthy:** Is somatosensory acuity linked to implicit motor adaptation? The latter is larger in old compared to younger people? In light of reliability-based sensory integration, we hypothesized that this larger implicit adaptation was linked to an age-related lower reliability of somatosensation. Over two experiments and 130 participants, we failed to find any evidence for this. We discuss alternative explanations for the increase in implicit adaptation with age and the validity of our somatosensory assessment.

## Introduction

As we grow older, our movements become less fluent, slower and more variable (Krampe, 2002; Yan et al., 1998). Besides altered movement parameters, aging also has an effect on the ability to adjust our movement to changes in the environment. The ability to adapt movement remains possible until old age. However, older adults are slower with respect to younger adults to adjust their movement and they do not always adjust their movement to the same extent as younger adults (Fernández-Ruiz et al., 2000; Seidler, 2007, 2006; Vandevoorde and Orban de Xivry, 2019). This age-related decline of motor adaptation is caused by the decline of the cognitive component of motor adaptation (Hegele and Heuer, 2010; Heuer and Hegele, 2014, 2008; Huang et al., 2017; Vandevoorde and Orban de Xivry, 2020, 2019). This is in contrast with the implicit component of adaptation, which appears to remain intact with aging (Heuer and Hegele, 2008; Vandevoorde and Orban de Xivry, 2019). The implicit component of motor adaptation, or internal model recalibration, is the adaptation of movement in response to sensory prediction errors (Morehead et al., 2017; Shadmehr et al., 2010; Taylor and Ivry, 2011). Sensory prediction errors occur when the actual sensory feedback is not matching with the predicted sensory feedback (Miall and Wolpert, 1996). For instance, in a typical visuomotor rotation experiment a rotation of the cursor is introduced with respect to the unseen hand movement: Participants expect visual feedback at the position of the hand, but it occurs at a different rotation angle. A sensory-prediction error causes a drift of the hand movement in the opposite direction of this error and this happens without conscious control (Morehead et al., 2017). Learning from sensory prediction errors appears to depend on the cerebellum since it is reduced in cerebellar patients (Morehead et al., 2017; Tseng et al., 2007). In a series of three different experiments, we recently demonstrated that implicit motor adaptation does not deteriorate with aging and sometimes even increased (Vandevoorde and Orban de Xivry, 2019). However, this observation of increased function is surprising since cerebellar brain structure shrinks with aging (Raz et al., 2005; Walhovd et al., 2011). In this paper, we want to test the hypothesis that age-related sensory deficits are linked to the increased reaction to sensory prediction errors. Indeed, the acuity of somatosensation (Goble et al., 2009; Shaffer and Harrison, 2007) declines with aging. For instance, human and animal studies have shown that the number and properties of cutaneous and joint mechanoreceptors are altered with age (Aydoǧ et al., 2006; Johnson, 2001; Miwa et al., 1995; Shaffer and Harrison, 2007; Swash and Fox, 1972; Verrillo et al., 2002). Furthermore, these age-related changes in the peripheral nervous system are accompanied by changes in the central nervous system that contribute to the decline in somatosensory function with aging (Goble et al., 2009).

One explanation for the increased internal model recalibration with aging is that age-related sensory deficits could lead to altered sensory integration, which might be based on the reliability of sensory inputs (van Beers et al., 2002). For instance, if we assume that age-related changes in visual acuity are smaller than those in somatosensory acuity, the weight of somatosensory input could decrease in elderly people compared to young people during optimal combination of visual and somatosensory inputs (Ernst and Banks, 2002). This up weighting of visual feedback could result in a larger estimated error, which would yield in turn an increased learning (learning rate is equal but estimated error is larger).

Another possibility is that age-related sensory deficits could lead to an increased reliance on the predictive pathway (Deravet et al., 2018; Kording and Wolpert, 2004; Orban de Xivry et al., 2013). Such a shift in the balance between somatosensory and predictive pathways might account for an increase in sensory attenuation with age (Parthasharathy et al., 2020; Wolpe et al., 2016). Sensory attenuation is the reduced perceived intensity of sensation from self-generated sensory feedback compared with external actions. Wolpe et al. (2016) observed a link between this increased sensory attenuation and altered somatosensory acuity with aging.

Following both of these explanations, our research hypothesis is that age-related somatosensory deficits are linked to the increased reaction to sensory-prediction errors (enhanced internal model recalibration).

Note that this hypothesis departs from other lines of research in which impairment in somatosensory function or disruption of the somatosensory cortex have been linked to an impairment in internal model recalibration. Some studies have suggested an impaired internal model for deafferented patients (Ghez et al., 1990, Vercher et al. 2003), while others suggested that deafferented patients can still trigger the internal model using alternative cues (Hermsdöfer et al., 2008). Recent work has shown that motor adaptation remains possible after proprioceptive loss (Miall et al., 2018) and that an increased weighting of vision vs proprioception was associated with force-field adaptation (Sexton et al., 2019). Furthermore, it has been shown that disruption of the somatosensory cortex impairs internal model recalibration (Mathis et al., 2017).

In order to verify our research hypothesis, we designed two paradigms that quantified internal model recalibration and upper limb somatosensory acuity, and investigated the relation between these two properties. In a first paradigm, we used a cued adaptation task that allowed us to assess explicit and implicit components of motor adaptation (Morehead et al., 2015). In a second (preregistered) paradigm, we assessed implicit adaptation by applying a task-irrelevant clamped feedback task in which participants’ reaching movement adapted implicitly to the visual error of constant size (Morehead et al., 2017). In both paradigms, we assessed somatosensory acuity in two ways: position sense, the ability to identify the static location of a body part, and kinesthetic sense, ability to identify body motion ((Goble et al., 2009; Herter et al., 2014)). We measured position sense with the arm position-matching task, which required participants to mirror-match positions of the right dominant arm with the left arm as accurate as possible without visual feedback (Dukelow et al., 2010). In a large cohort of 209 individuals, it is shown that this ability declines with aging (Herter et al., 2014). We measured kinesthetic sense with a custom-developed task that resembled a previously described perceptual boundary tasks (Darainy et al., 2013; Kitchen and Miall, 2020; Ostry et al., 2010). Our research hypothesis predicted a negative relationship between internal model recalibration and somatosensory acuity. That is, the more somatosensory acuity is degraded in an individual, the larger the recalibration of the internal model is.

## Methods

### Participants

In total 133 healthy adults were recruited and participated after providing written informed consent. Sixty-two participated in paradigm 1 and 71 participated in paradigm 2.

Sixty-two participants were included in the final analyses for paradigm 1 (Table 1). These 62 participants consisted of 30 young adults (between 20 and 32 years old, age: 22.9 ± 2.7 years, mean ± SD; 15 females) and 32 older adults (between 61 and 75 years old, age: 67.6 ± 4.5 years; 13 females). The somatosensation data from these participants has not been presented before, while the adaptation result were already presented in (Vandevoorde and Orban de Xivry, 2020) (as dual-task design E2).

**Table 1:**
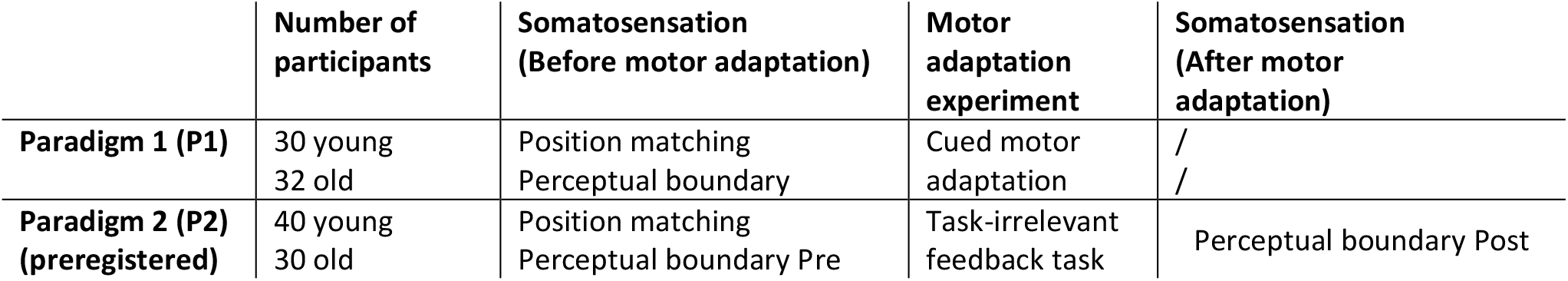
Description of the two paradigms for assessing somatosensory acuity and internal model recalibration.

|  | Number of participants | Somatosensation (Before motor adaptation) | Motor adaptation experiment | Somatosensation (After motor adaptation) |
| --- | --- | --- | --- | --- |
| Paradigm 1 (P1) | 30 young<br>32 old | Position matching<br>Perceptual boundary | Cued motor adaptation | /<br>/ |
| Paradigm 2 (P2) (preregistered) | 40 young<br>30 old | Position matching<br>Perceptual boundary Pre | Task-irrelevant feedback task | Perceptual boundary Post |

Seventy participants were included in the final analyses for paradigm 2 (Table 1). Data from one older adult were excluded because she didn’t move with the right speed to the targets and often started reaching movements before the targets appeared. The 70 participants consisted of 40 young adults (between 19 and 30 years old, age: 22.0 ± 2.5 years, mean ± SD; 16 females) and 30 older adults (between 61 and 75 years old, age: 67.3 ± 4.1 years, mean ± SD; 10 females).

The Edinburgh handedness questionnaire (Oldfield, 1971) revealed that all participants were right-handed. All participants were screened with a general health and consumption habits questionnaire. None of them reported a history of neurological disease or was taking psychoactive medication. In older adults general cognitive functions was assessed using the Mini-Mental State Examination (MMSE) (Folstein et al., 1975). All elderly scored within normal limits (MMSE-score ≥ 26). The protocol was approved by the local ethical committee of KU Leuven, Belgium (project number: S58084). Participants were financially compensated for participation (10 €/h).

### Experimental setup

For somatosensory acuity testing (Figure 1A-C), we used the KINARM End-Point Lab robot (BKIN Technologies Ltd., Kingston, Canada). Participants were seated in a chair and controlled the handle of the right robot with their right dominant hand. During the two somatosensory tasks, visual feedback of their own hand was blocked. Visual feedback was displayed on the virtual reality display component of the End-Point Lab on a semi-transparent mirror, horizontally mounted in front of the participant. This feedback was provided as a white cursor dot, which was properly aligned with the real position of their hand. The End-Point Lab allowed real-time control and data acquisition at 1 KHz.

**Figure 1:**
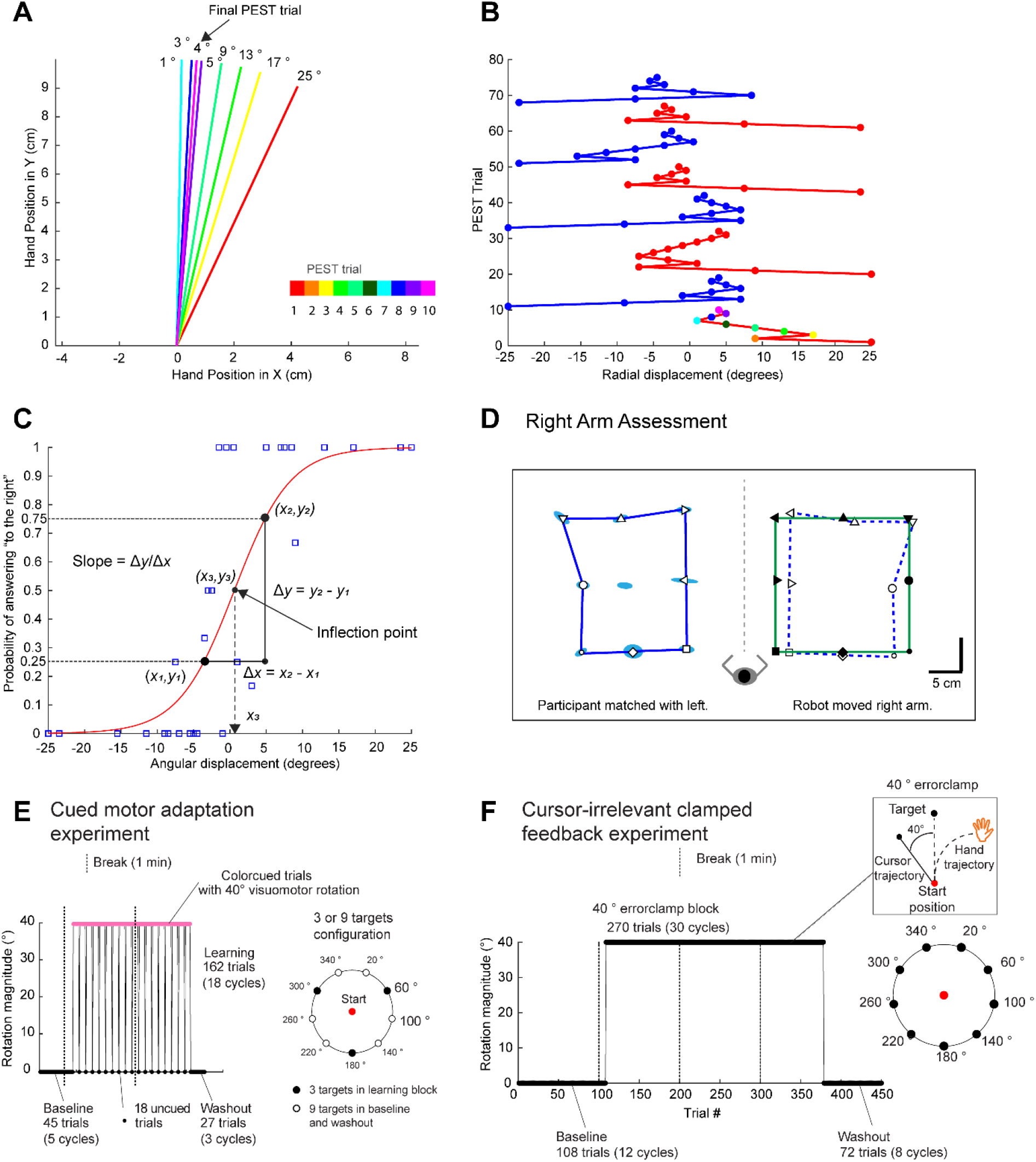
Experimental paradigm for assessing somatosensory acuity and internal model recalibration. **A)** Perceptual boundary test. Experiment is designed to test how well participants can discriminate angular deviations. An example of first PEST run for with an initial deviation of 25 °. **B)** An example of a sequence of eight PEST runs for one (imaginary) participant. The first run is the same as the one shown in Fig 1A. The angular direction is shown on the horizontal axis and the PEST trial number shown on the vertical axis. The colored sequence is the same as the sequence indicated in fig1A. **C)** Psychometric accuracy curve with indication of inflection point and slope. Y-axis shows the probability of answering “deviation to the right” for a specific amount of angular displacement. Psychometric curve is constructed with data from Fig1B. **D)** Right arm assessment of position matching task for one (imaginary) participant. Robot moved the right arm and participant matched with the left arm. **E)** Cued motor adaptation experiment. **F)** Cursor-irrelevant clamped feedback experiment. The experiment background for the visuomotor rotation experiment was black and targets were white, the opposite as visualized here in Figure 1E-F.

For visuomotor rotation experiments (Figure 1E-F), participants were seated in front of a table. With their right hand, participants performed a reaching task on a digitizing tablet (Intuos pro 4; Wacom Co. Ltd., Kazo, Japan) with a digitizing pen. A wooden cover above the tablet prevented visual feedback from their moving arm. Movement trajectories were recorded at 144 Hz. Visual feedback was displayed on a 27 inch, 2560 x 1440 optimal pixel resolution LCD monitor with 144 Hz refresh rate (S2716DG, Dell), vertically mounted in front of the participant.

### Experimental protocols

#### Organization of experiments

Two different paradigms were used (Table 1). Paradigm 1 consisted of an assessment of somatosensation before the cued motor adaptation experiment. Paradigm 2 consisted of an assessment of position matching and perceptual boundary before the task-irrelevant feedback task, which was followed by a second assessment of perceptual boundary only.

#### Preregistration

Design and analysis procedure of Paradigm 2 were preregistered online: https://osf.io/qg3t2/registrations. This preregistration included the main hypotheses, the key dependent variables, the amount of participants, the main analyses and some of the secondary analyses investigated. In the preregistration, we mentioned that we would include 30 participants per group. However, for this study, we included 10 additional subjects in the young adults group but these additional participants did not change the outcome of our study. Including these 10 additional subjects or not, did not yield different results for the task-irrelevant feedback experiment (Figure 5), for the arm position matching task and perceptual boundary test (Figure 6) or for the relationship between adaptation and somatosensation (Figure 7). To be concise, we reported only the results with all participants included.

#### Perceptual boundary test

The perceptual boundary test (Figure 1A-B) was an adapted version of the test described in (Ostry et al., 2010). In our task, participants made right arm reaching movement while their movement was constrained to a certain angular deviation relative to a straight line to the target. No visual feedback was provided during reaching; only at the end of the return to the center, a short instance of visual feedback was applied to simplify the return to the start position. Returning to start was simplified by using a viscosity gradient. Participants were instructed to verbally indicate whether they were deviated to the left or to the right of a reference position (straight ahead). By gradually decreasing the angular deviation, it became possible to estimate how accurately participants could discriminate angular deviations. In order to obtain an efficient estimation of the perceptual boundary for each individual, a parameter estimation by sequential testing (PEST) procedure was applied (Taylor and Creelman, 1967). This algorithm starts with a large deviation and depending on the individual’s response, it decreases the size of the deviation progressively until the deviation falls below a minimum threshold. Every time an individual uses the same response for the current trial compared to the previous trial (i.e. “right” stays “right”, or “left” stays “left”), the size and direction of the angular deviation in the next trial remain the same as for the current trial. Every time an individual uses a different response for the current trial compared to the previous trial (i.e. “right” becomes “left”, or “left” becomes “right”), the step size of the angular deviation will halve and the deviation will switch direction in the next trial compared to the current trial. In our task, eight PEST runs were applied. The first four started with an initial angular deviation of 25 °, the last four started with and initial angular deviation of 23.5 °. The first initial deviation was always 25 ° to the right of the target (Figure 1A). Every following PEST run had a first initial deviation with an opposite direction from the previous PEST run initial deviation. For example, the first PEST run was 25° to the right and the second PEST run was 25° to the left. In total four PEST runs started from the right and four started from the left. Our minimum threshold was 1.5 °: As soon as a step size fell below this minimum threshold, the current PEST would stop and the next PEST run would be initiated.

In the original task of (Ostry et al., 2010), lateral deviations were used instead of angular deviations. We induced angular deviations in order to make the boundary test more similar to the deviations experienced during visuomotor rotation. Also other parameters of the task were resembling our visuomotor rotation experiment: the reaching distance from start to target was 10 cm and reach timing was between 125 ms and 375 ms. However, only one target was presented in front of the participant, instead of nine targets as in the visuomotor rotation experiments.

First, participants could get familiar with the required reaching speed. They received the following instructions: “First, you can practice the speed of the movement and afterwards I will explain the real task. You have to move a white dot to the red square in the center by moving your right arm. As you enter the red square, a white circle of 10 cm radius will appear and the white dot will disappear. A small gap exists at the top of the circle, you will have to move your hand through this gap with the right speed. If you move too slow, the circle will become blue. If you move too fast, the circle will turn orange. If you move with the correct speed, the circle becomes green.” Participants could practice the speed of their movement until they moved a couple of times with the correct speed. After this, we started perceptual boundary test and instructed participants: “Now, we start the real task. In this task, you will make reaching movements with the same speed as before. However, now the robot will constrain your movement to a certain direction, either to the left of the gap or to the right of the gap. Your task is to say to me whether you think it was to the left or to the right. In every trial a deviation exists, so you always have to say either left or right, you can also guess if you are not sure.”

#### Arm Position Matching task

The Arm Position Matching task was available as a Standard Test on the Dexterit-E Software provided with the KINARM End-Point Lab. The test allowed assessing position sense of the hands in horizontal workspace (Dexterit-E 3.7 User Guide). In this test, the robot passively moved one hand to a particular position and participants were required to mirror-match this position with their other hand (Figure 1D). This requires passive position sense for one arm and active movement and position sense of the other arm (Dukelow et al., 2010).

In this task, the robot passively displaced the right hand and then stopped at a particular location. The participants had to match the position of the right hand with their left hand in a mirror fashion. There were nine possible targets organized in a 3×3 matrix. Participants could take as much time as they wanted and had to indicate verbally when they had matched the position. After the verbal response, the examiner started the following trial. Each of the nine target locations was randomly presented in one block. In total, six blocks or 54 trials were assessed for each participant.

#### Cued motor adaptation

This visuomotor rotation experiment was described before as experiment E2 in (Vandevoorde and Orban de Xivry, 2020), here it is part of our first paradigm (Table 1). The cued motor adaptation experiment allowed to assess implicit and explicit adaptation by the introduction of a color cued cursor that indicated the presence or absence of a 40° visuomotor rotation (Figure 1E). A short reaching baseline of 45 trials (5 cycles) was implemented. A learning block of 162 trials (18 cycles) and a short washout of 27 trials (3 cycles) followed the baseline. A single start point location (filled red circle) was used. Nine targets (open and filled white circles on black background) were presented during reaching trials of the dual-task baseline and during baseline and washout trials. Three targets (filled white circles on black background) were used during the learning block. Nine targets were presented pseudo-randomly in cycles during baseline and washout with each of the nine targets presented once per cycle. In the learning block, 9-trial-cycles consisted of three 3-trial-subcycles because only three targets were used. In each subcycle, each of the three targets was presented once. The learning block consisted of 18 cycles (or 54 subcycles or 162 trials).

The cursor dot remained white the entire baseline and washout blocks. However, during the two adaptation blocks, the cursor became a pink square (i.e. cued trial) instead of a white cursor dot. This cue indicated the presence of a 40° rotation. In each adaptation block, the cursor became again a white cursor dot (i.e. uncued trials) for a few trials, indicating the absence of the perturbation. The instructions were: “First, the cursor will be a white dot, but at some point the cursor will change to a pink square. At that moment, something special will happen but you still have to try to do the same thing, reach to the target with the cursor. The cursor will sometimes change back to a white dot. These trials with a white dot are normal reaching trials like in baseline.” The change in behavior induced by the cue was thus a measure of the explicit component of adaptation as participants could use the cue to switch off any conscious strategies they were applying to counteract the perturbation (Morehead et al., 2015). We reinforced the awareness of cue switches (signaling a cued trial among uncued ones or an uncued trial among cued ones) with a warning sound played before each cue switch and with a text that indicated the cue switch, displayed for 5 s: 'Attention! The color of the cursor has changed.' Eighteen uncued trial were presented in the adaptation block. These uncued trials were equally distributed over the whole adaptation block and among the three targets (six uncued trials per target).

One trial of the cued adaptation experiment took exactly 4.5 seconds. First, participants performed a target reach. Immediately after the reaching, extra fixation time was added in order to obtain exactly 4.5 seconds per trial. After 4.5 seconds, the next trial started automatically. If participants exceeded 3 seconds for the reaching movement, a warning sign was shown and a high pitch was played in order to instruct participants to speed up. The correct reaching time was between 175 ms and 375 ms, which was indicated with a target color change. If reaching time was above 375 ms, the target became purple. If reaching time was below 175 ms, the target became red. Two breaks of 1 min were given to participants, one before the 4^th^ cycle (trial 36) of the baseline and one before the 9^th^ cycle (trial 81) of the learning block.

#### Task-irrelevant clamped feedback

The task-irrelevant clamped feedback experiment assessed internal model recalibration and is adapted from Morehead et al. (2017). This experiment was part of our preregistered second paradigm (Table 1). In this visuomotor rotation task, the cursor direction of motion was made completely irrelevant by dissociating it from the hand direction of motion (Figure 1F). That is, in these task-irrelevant clamped feedback trials, participants were instructed to ignore the cursor that is always rotated 40 ° with respect to the target direction and to try to move their hand accurately towards the target in the absence of relevant visual feedback of hand position. Targets were presented in cycles of nine trials with each cycle consisting of the nine targets presented randomly (Figure 1F). In total, the experiment consisted of 450 trials or 50 cycles: Baseline consisted of 12 cycles (or 108 trials), task-irrelevant clamped feedback trials were presented for 30 cycles (or 270 trials) and washout consisted of eight cycles (or 72 trials). One-minute breaks were given to the participants before trial 99, 200 and 300. In baseline, participants could win points for accuracy. However, during the adaptation block, participants were clearly informed that it was not possible to win extra points, because the cursor could never reach the target during these trials. To keep participants motivated during the adaptation block, the amount of remaining trials was visualized on the monitor. Every trial had a fixed duration of 2.5 s/trial. If the reach and return to the start location were performed in less than 2.5 s, an extra amount of time was added to obtain exactly 2.5 s for the current trial before starting the next trial. During this extra amount of time, we only showed the cursor dot and the start position. If taking longer than 2.5 s for the reaching or for the reaching & returning, the next trial was automatically initiated. The trial duration was rather long, because we wanted to make sure that young and older adults behaved similarly during target reaching. Older adults, in general, took longer to return back to the start position. The reaching time to the target was constrained to a minimum of 125 ms and a maximum of 375 ms. If reaching time was below 125 ms the target become red and low-beep sound played, if above 375 ms the target become purple and a high-beep sound played. During cursor-irrelevant feedback trials, only the sound feedback indicated incorrect reaching time. Participants could get familiar with the experiment by practicing 10 baseline trials, 10 cursor-irrelevant feedback trials (of different rotation directions) and 10 washout trials.

#### Additional details for the two visuomotor rotation experiments

Each visuomotor rotation experiment consisted of a series of reaching movements to a single white circular target located 10 cm away from the central starting position. For each trial, the participant had to rapidly move his or her right hand to move a white cursor through the target.

The feedback cursor, which represented hand position (when there was no perturbation) was visible until movement amplitude exceeded 10 cm. At this point, a white square marked the position where movement amplitude reached 10 cm, providing visual feedback about the end point accuracy of the reach. The white square had sides of 5 mm. The cursor position froze at the end of each reaching movement and was visible for 1 s. The two visuomotor rotation experiments first started with baseline trials (no perturbation) with normal cursor feedback (i.e. the cursor represents the actual hand position) and continued with perturbation trials where the feedback was either rotated or irrelevant. Before baseline, each participant performed some familiarization trials to make sure that they understood the instructions and that they performed the task correctly.

The diameters of the starting point and the target were both 10 mm and the cursor dot had a diameter of 5 mm. Targets were presented in pseudo random order in cycles of nine trials per cycle. Target position are shown in Figure 1E-F.

While returning to the central starting position, the cursor disappeared and only a half white arc of 180° (i.e. return arc) was visible. The radius of the return arc depended on the position of the pen on the tablet, i.e. the radius of the arc was equal to the angular distance between the position of the hand and the starting point. The center of the return arc was the central starting position. To reduce the time for returning to the starting point, the cursor became visible as soon as the hand was within a region near the central starting position (20 mm x 20 mm). The reach area was divided in three different zones of 120 °. The return arc was centered in the same 120° zone as where the participant’s (invisible) hand was. Participants had to move their hand in the opposite direction of the arc in order to return to the starting location. The arc allowed participants to return to the starting position and at the same time prevented the participants from using the visual feedback during the return movement to learn about the perturbation. If taking longer than 1.5 s to return, the cursor became visible again as such returning to start became very easy.

Participants were instructed to score as many points as possible by hitting the target with the cursor. When hitting the target, the participant received 50 points. When hitting targets correctly on consecutive trials, 10 bonus points were received for every additional trial with a correct hit (e.g. 60 points were received the second trial after hitting targets correctly on two consecutive trials, 70 points were received the third trial after hitting targets correctly on three consecutive trials). When reaching in close proximity of the target, the participant received 25 points. In P2, the zone for receiving 25 points was an additional 6 mm at both sides of the target. In P2, the zone for receiving 25 points was an additional 5 mm at both sides of the target. The reward for near misses is implemented for keeping participants motivated even when they are not achieving very high accuracies. This was clearly instructed on beforehand. To receive points, participants were required to reach the target between 175 and 375ms after movement onset. The cumulative score of all previous trials was displayed throughout the experiment, except during the cursor-irrelevant feedback block. During cursor-irrelevant feedback trials, no score could be obtained since the cursor could never hit the target.

The experimenter (KV) was present during the entire experiment to motivate the participants to achieve the highest possible score and to make sure that the participant performed the task correctly. The experimenter regularly reported that the participant was performing well, even when the score was below average.

### Data analysis

#### Data availability

Data and scripts will be available on Open Science Framework (OSF) (https://osf.io/qg3t2/?view_only=30b338a6792c45c2b229838a7f2a216e).

#### Preprocessing

In task-irrelevant feedback experiment (P2), the first trial after each of the two breaks of the adaptation block was removed since only one older participant could respond quick enough. The other older adults were not yet ready to make a reaching movement within the 2.5 s trial time after the 1 min break. Removing this one trial did not affect our comparison between young and older adults.

#### General

The statistically significant threshold was set at p<0.05 for the ANOVA’s, t-tests and robust linear regression. We reported effect sizes (eta squared *ƞ*^2^ for ANOVA and Cohen’s d for two-sample t-test) as well as F or t and p-values. For robust linear regression, we reported standardized beta coefficients β and p-values of these coefficients. Multiple comparisons are corrected by applying Bonferroni corrections. Confidence intervals (CI) are reported for eta squared *ƞ*^2^, Cohen’s d and standardized beta coefficients β.

For eta squared *ƞ*^2^, we calculated CI 95 % by 10.000 bootstraps of the two age groups and calculating the corresponding statistics. For Cohen’s d, CI 95 % was calculated with the formula:

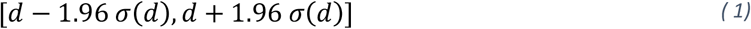

With *σ*(*d*):

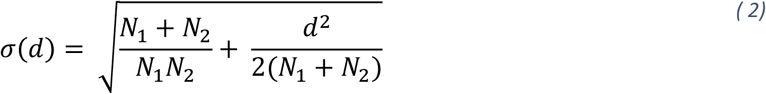

as specified in Hedge and Olkin (2014). For standardized beta coefficients β, we used the following formula:

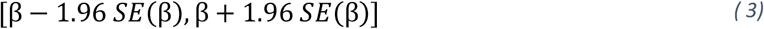

With SE(β), the standard error of the standardized beta coefficients β.

#### Analysis 1: Implicit adaptation level

In paradigm 1, implicit adaptation was assessed with cued motor adaptation. We applied the same analysis as described in (Vandevoorde and Orban de Xivry, 2020). The adaptation block of cued adaptation was corrected for baseline errors by subtracting the average error of the last two baseline cycles. We analyzed the data in all the uncued trials of the learning block (18 uncued trials). The amount of implicit adaptation was calculated as the average of the uncued trials (Morehead et al., 2015). One 2-way ANOVA was used, with the between-subject factors, age and rotation direction, and with the implicit adaptation as dependent variable.

In task-irrelevant clamped feedback task (preregistered paradigm 2), the hand angle of the last ten cycles (90 trials) of the clamped visual feedback block were averaged to obtain the implicit adaptation level. Baseline accuracy measurement of unperturbed reaching allows to control for age-differences in unperturbed conditions. All adaptation trials are corrected for baseline accuracy by subtracting the average error of the last two baseline cycles (18 trials). One 2-way ANOVA was used, with the between-subject factors, age and rotation direction, and with the implicit adaptation as dependent variable.

#### Analysis 2: Position matching task

This analysis was **preregistered**for paradigm 2. Somatosensory acuity was defined as the absolute error XY and as the variability XY (Dukelow et al., 2010; Herter et al., 2014). For calculating variability, first the standard deviation was calculated for each of the six target locations for x and y direction separately. Next, the mean of these standard deviations of all six target locations resulted in the variability for the x and y direction (*Var_x_*, *Var_y_*). Variability XY was calculated with the following formula:

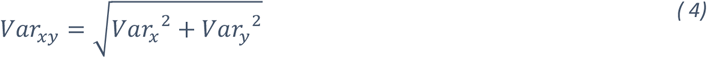

For calculating absolute error (systematic shift) between the active and passive hand the same procedure was followed. First, absolute error was calculated for each of the six target locations for x and y direction separately. Next, the mean of these absolute errors of all six target locations resulted in the absolute error for the x and y direction (*AbsError_x_*,*AbsError_y_*). Absolute errors XY was finally calculated as follows:

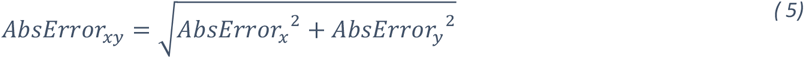

A separate unpaired two-tailed t-test was used to compare each variable (*Var_xy_* and *AdsError_xy_*) between the two age groups.

#### Analysis 3: Slope and inflection point from perceptual boundary test

This analysis was **preregistered** for paradigm 2. Each trial of the perceptual boundary test resulted in a binary response, either left or right. The probability of a “rightward direction” answer could be calculated from the responses that participants gave for all deviations that they experienced. This allowed us to create a sigmoidal probability response curve (Figure 1C) as a function of experienced angular deviations. Function, *glmfit,* was used in Matlab to fit the probability of a “rightward direction” answer to the tested angular deviations with a logistic regression function, with p, the probability, b, the coefficient estimate, X, the tested angular deviations:

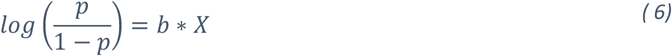

The output of this function was the coefficient estimate b. Afterwards, we provided the coefficient estimate, b, to the *glmval* function, which computed predicted values for all angular deviations that were not tested between −25 ° and + 25 °. The final output was the psychometric curve for each participant that showed the probability of answering “right” for every angular deviation between – 25 ° and + 25 °.

The psychometric curve allowed us to calculate the following two preregistered variables: The first variable was the slope of the psychometric accuracy curve, measured as the slope of the straight line between the 25 % and the 75% chance points. The second variable was the inflection point of the psychometric curve, more specifically the point that crossed the x-axis (x3 in Figure 1C). The slope describes how sensitive someone can discriminate angular displacements during reaching movements. The inflection point describes which direction someone feels as the zero angular displacement, i.e. straight ahead of the body of the participant.

The perceptual boundary test was performed before the cued adaptation experiment (P1), which resulted in two output variables: slope and inflection point. An unpaired two-tailed t-test was used to compare each variable between the two age groups. In P2, the perceptual boundary test was also performed before and after the task-irrelevant feedback experiment, which resulted in four output variables: slope-pre, slope-post, inflection point-pre and inflection point-post. A repeated-measures ANOVA was used, with the within-subject factor, test time, with the between-subject factors, age and rotation direction, and with slope as dependent variable. A second repeated-measures ANOVA was performed, with the same within-and between-subject factors, but with inflection point as dependent variable.

#### Analysis 4: Relationship between implicit adaptation level and somatosensation (Paradigm 1)

Robust linear regression (robustfit in Matlab) (Holland and Welsch, 1977) was performed between the measure of implicit adaptation and somatosensory acuity in order to verify that correlations between these two variables were not influenced by between group differences in the variables. Implicit adaptation (Y) was estimated using a linear combination of somatosensation variable (X), a binary age vector (G) and the interaction of X and G in the regression equation with intercept A and regression coefficients (B,C,D):

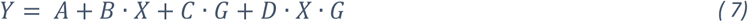

Standardized beta coefficients (β) will be obtained instead of regression coefficients (B,C,D) when first converting variables X and Y to z-scores and afterwards applying linear regression.

We applied robust linear regression (Equation (7)) four times with data from paradigm 1 and corrected for multiple testing with Bonferonni (*α* = 0.0125). For every regression, the implicit adaptation level as described in Analysis 1 was used as the Y-variable. The X-variable could be any of the two measures of somatosensation as described in Analysis 2 (Var XY and AbsError XY), or any of the two measures of somatosensation as described in Analysis 3 (slope, inflection point).

#### Analysis 5: Preregistered relationship between implicit adaptation level and somatosensation (Paradigm 2)

In addition, for paradigm 2, robust linear regression (Equation (7)) was a preregistered analysis to investigate the hypothesis that increased internal model recalibration was related to somatosensation decline. In total, robust linear regression (Equation ( 7)) was performed four times for paradigm 2. We corrected for multiple testing with Bonferonni (*α* = 0.0125). For every regression, task-irrelevant implicit adaptation level as described in Analysis 1 was used as the Y-variable. The X-variable could be any of the two measures of somatosensation as described in Analysis 2 (Var XY and AbsError XY), or any of the two measures of somatosensation as described in Analysis 3 (slope-pre, inflection point pre).

#### Analysis 6: Relationship between implicit adaptation and change in somatosensation (Paradigm 2)

We calculated the change in inflection point between post and pre condition. This difference was multiplied with −1 for participants that experienced a clockwise rotation in paradigm 2 in order to take into account the influence of perturbation direction. This difference was compared to zero with a one-sample t-test.

Robust linear regression was used to verify whether a relationship existed between difference of inflection point (Analysis 4) and level of internal model recalibration (Analysis 1). Additionally, we verified whether the aftereffect at the end of the washout period of task-irrelevant feedback experiment (P2) was correlated with the difference of inflection point.

#### Analysis 7: Comparison between somatosensory acuity from position matching task and perceptual boundary test

Data were obtained for both tests in both paradigms (P1 & P2), which allowed us to combine data of all 133 participants for this comparison. Four variables were described for the position matching task (Analysis 2: Var XY and AbsError XY) and two variables were described in the perceptual boundary test (Analysis 3: slope, inflection point). We didn’t include the measurement of slope and inflection point from the post-condition since this was only available for paradigm 2. Robust linear regression as described in Analysis 4 was used to compare the variables from both somatosensory acuity tests. Since the two variables of position matching should be compared with the two variables of perceptual boundary test, we had to perform the robust linear regression four times. We corrected for multiple testing with Bonferonni (*α* = 0.0125).

## Results

We investigated whether an age-related change in internal model recalibration was related to a decline in somatosensation with aging. According to our **preregistered hypothesis**, somatosensory acuity would decline with aging and this age-related decline in somatosensory acuity would be related to the age-related increase of internal model recalibration. To this end, we assessed internal model recalibration with a visuomotor rotation experiment and tested the relationship between the level of internal model recalibration and somatosensory acuity. We quantified somatosensation of the upper limb via a position matching task and a perceptual boundary test that provided us with a measure of, respectively, position and kinesthetic sense.

### No evidence that somatosensory acuity is related to implicit adaptation

First, we quantified internal model recalibration via implicit adaptation during a cued motor adaptation experiment (Paradigm 1 (P1) Table 1; Figure 1E, (Morehead et al., 2015)) for 30 young (age range: 20-32 years old) and 32 older (age range: 61-75 years old) participants. During the cued motor adaptation experiment, color cues allowed participants to switch their explicit aiming on or off (Figure 1E). The level of implicit adaptation was similar between young and old (Analysis 1) (Figure 2A-B; F(1,58) = 1.6, p = 0.2, η^2^ = 0.02, CI95-η^2^ = [0.0000; 0.1179]). Cued adaptation results were described before in (Vandevoorde and Orban de Xivry, 2020), we show these results again for completeness. We observed no decline of implicit adaptation with age as measured with cued motor adaptation (Figure 2).

**Figure 2:**
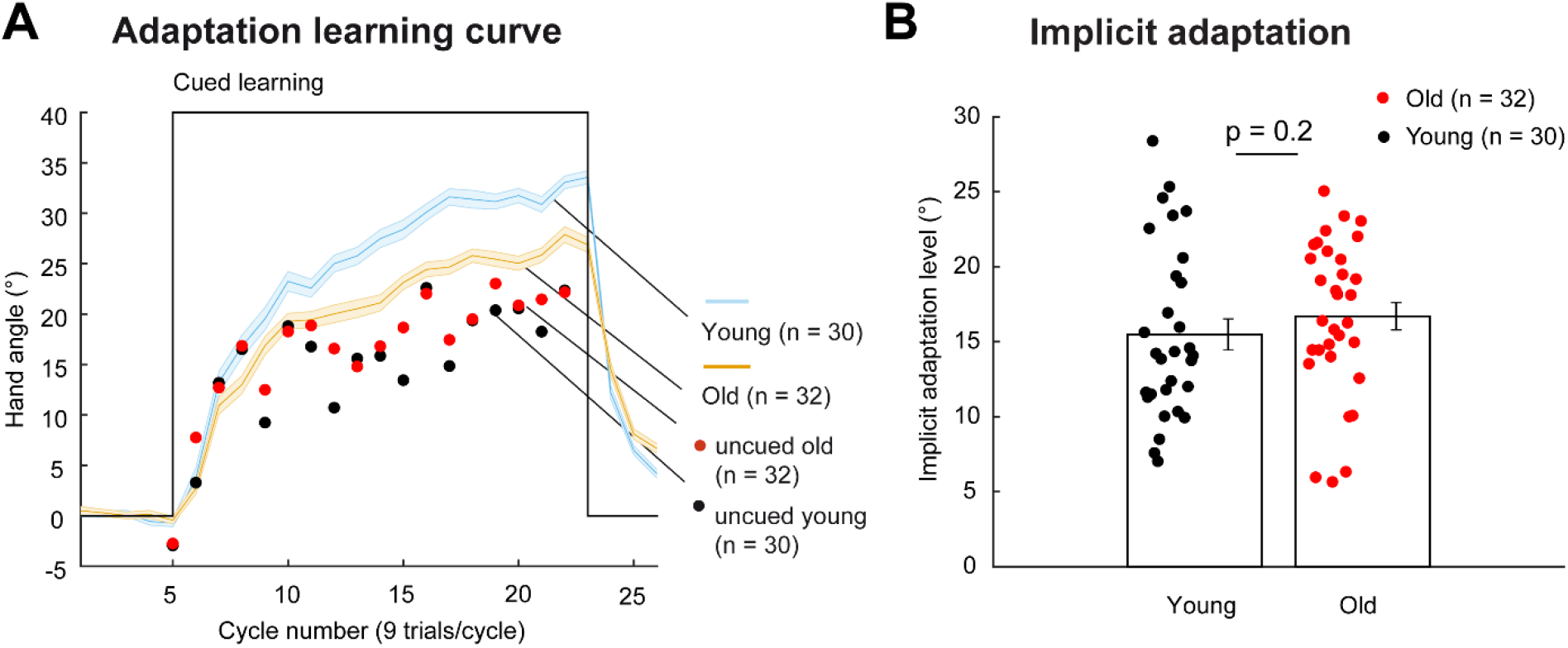
Results of cued motor adaptation. **A)** Evolution of the hand angle over the course of the experiment during cued (blue solid lines for young participants and orange ones for old participants) and uncued trials (red dots for older adults and black dots for younger adults). Each dot shows the hand angle during one uncued trial as the average of the whole group (young or old). The area around the solid lines represents the standard error of the mean. **B)** Same data as in Figure 2A, but shown as individual levels of implicit adaptation measured during uncued trials. Each dot represents the data of one participant, which is the average of the hand angles during all 18 uncued trials (dots in panel A). The height of the bars represents the inter-individual means and the error bars represent the standard error of the mean.

We measured somatosensory acuity for right dominant arm position and arm movement with two different methods. The first method was the position matching task, in which a robot passively moved the right hand to a position and participants were required to mirror-match this position with their left hand (Dukelow et al., 2010). This provided a measurement of position sense. The second method was the perceptual boundary test to assess kinesthetic sense (Figure 1A-C). In the position-matching task, we focused on two variables, absolute error XY and variability XY (Equations (4) (5); Analysis 2). We did not find any evidence that performance for position-matching was reduced with aging in paradigm 1 (Figure 3A-B; Analysis 2: VarXY: t(60) = 0.65, p = 0.5, d = 0.17, CI95-d = [- 0.33, 0.66]).; AbsErXY: t(60) = −0.08, p = 0.9, d = - 0.02, CI95-d = [- 0.52, 0.48]). In addition, in the perceptual boundary test, we defined two additional variables, the slope and the inflection point of the psychometric accuracy curve (Analysis 3). The perceptual boundary test did not provide any evidence that the slope and the inflection point of the psychometric accuracy curve were different for young and old (Figure 3C-D; Analysis 3: slope: t(57) = −1.67, p = 0.1, d = - 0.43, CI95-d = [- 0.94, 0.07]; inflection point: t(57) = 0.03, p = 0.98, d = 0.007, CI95-d = [- 0.49, 0.51]).

**Figure 3:**
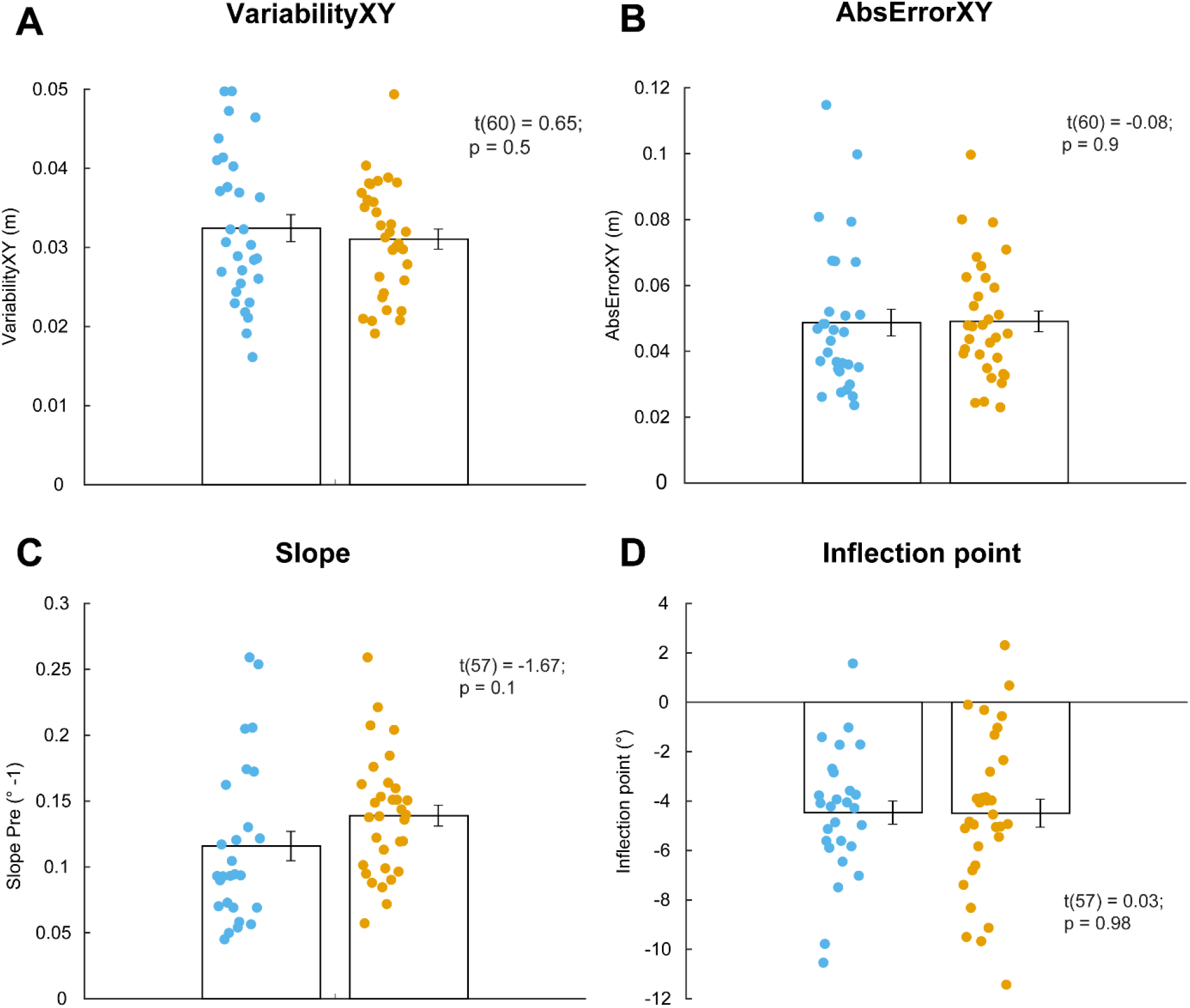
Outcomes of the somatosensory tasks, visualized with data from participants of paradigm 1. **A-B:** Outcomes of the position matching task measured before adaptation: variability of hand position, VariabilityXY, and the absolute error, AbsErrXY. Lower values are associated with better somatosensory acuity. **C-D:** Outcomes of the perceptual boundary estimation task measured before adaptation. For panel C, higher values are associated with better somatosensory acuity. No such interpretation exists for the inflection point. Each dot represents the data from one participant. The height of the bars represents the inter-individual means and the error bars represent the standard error of the mean. Three of the 30 young participants did not execute the perceptual boundary test.

After verifying how aging affects the level of implicit adaptation and somatosensory acuity, we explored the relation between these two properties. A decline in somatosensation was expected to be related to an increased internal model recalibration (negative relationship). However, we failed to find evidence that the position matching variables was related to the level of implicit adaptation (Equation (7); Figure 4A-B; Analysis 4; Table 2; β_VAR-XY_ = 0.11; p = 0.4, R = - 0.04; β_ABS-XY_ = - 0.14; p = 0.3, R = - 0.16). Furthermore, we could not find evidence in favor of a relationship between the perceptual boundary variables and implicit adaptation level (Figure 4C-D; Analysis 5; Table 2; β_SLOPE_ = - 0.18; p = 0.2, R = - 0.13; β_IP_ = - 0.04; p = 0.8, R = - 0.05). Since paradigm 1 did not provide evidence for the hypothesis, we wanted to confirm the absence of relationship in a second cohort of participants (Table 1). In this second paradigm, we assessed internal model recalibration with the task-irrelevant feedback experiment (Figure 1), which allows assessing internal model recalibration independently of the presence of explicit strategy. In addition, the task-irrelevant feedback experiment resulted in an increased level of internal model recalibration in our previous work (Vandevoorde and Orban de Xivry, 2019).

**Figure 4:**
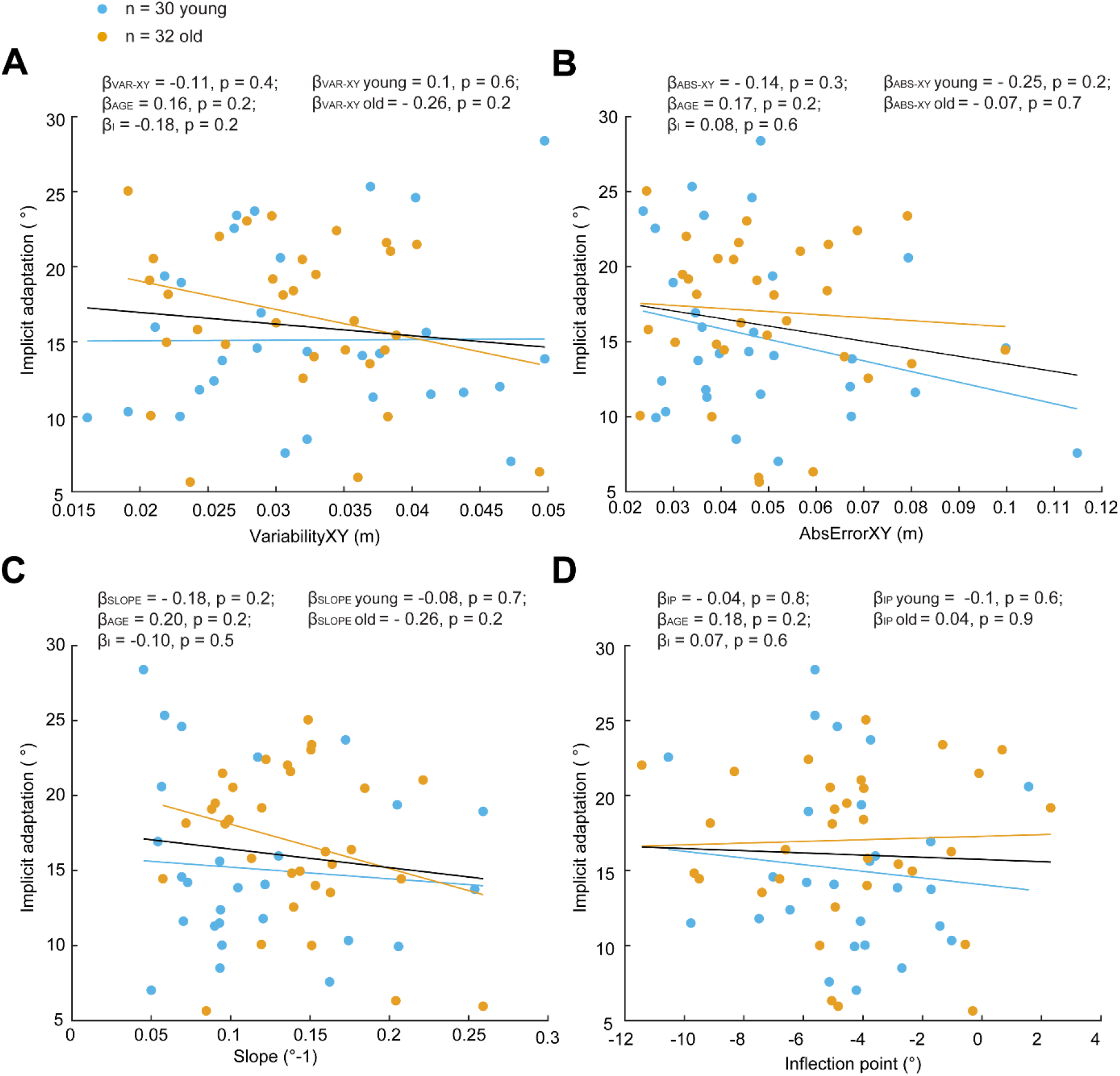
Relationship between implicit adaptation and somatosensory acuity. Data from paradigm 1. **A-B**: Scatterplot of the outcomes of the position matching task (data from fig.3A and B) and the extent of implicit motor adaptation from the cued motor adaptation task (data from Fig.2B). A positive relationship is expected given that better somatosensory acuity is associated with lower values. **C-D**: Scatterplot of the outcomes of the perceptual boundary task (data from fig.3C and D) and the extent of implicit motor adaptation from the cued motor adaptation task (data from Fig.2B). A negative relationship is expected for panel C given that better somatosensory acuity is associated with higher slope values. In all panels, each dot represent the data of a single participant. Solid lines represent the regression line from the robust regression analysis for both populations together (black line), for young participants only (blue lines) and for old participants only (orange lines).

### Evidence for an age-related decline in somatosensory acuity but no evidence that this is related to the age-related increase in internal model recalibration

In the second **preregistered** paradigm (Table 1), we assessed internal model recalibration by applying task-irrelevant clamped feedback in 40 young (age range: 19-30 years) and 30 older (age range: 61-75 years) participants. The motion of the cursor away from the target elicited drift in hand position (Figure 1E,(Morehead et al., 2017)). The change in movement direction was larger for older adults compared to younger adults (Analysis 1)(Figure 5A-B; F(1,66) = 8.5; p = 0.005; η^2^ = 0.09; CI95-η^2^ = [0.0065; 0.2834]). Two older participants did not follow instructions well as indicated by negative hand angles (Figure 5B), if we would remove these participants as outliers, the effect would be even stronger (CI95-η^2^ = [0.0726; 0.3617]).

**Figure 5:**
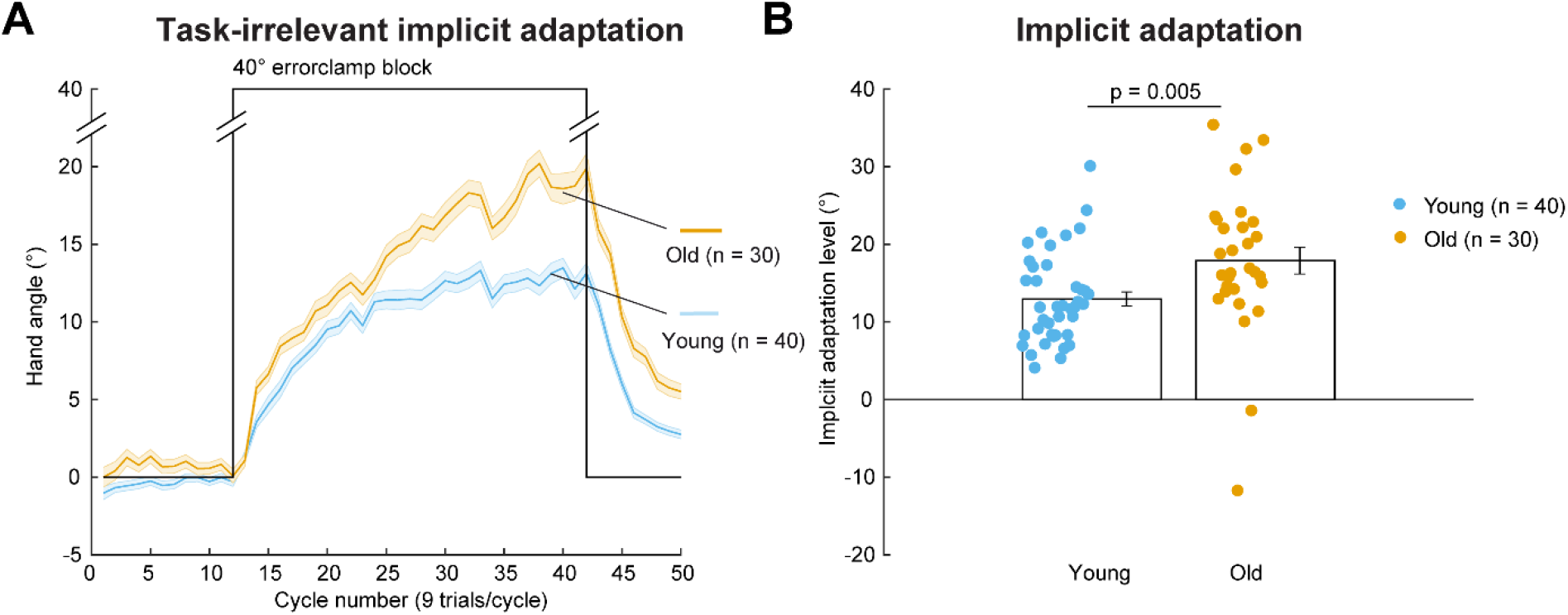
Results of task-irrelevant clamped feedback experiment for young and old: A) Evolution of the hand angle over the course of the experiment (blue solid lines for young participants and orange ones for old participants). The area around the solid lines represents the standard error of the mean. B) Same data as in panel A, but shown as individual levels of implicit adaptation measured during the last 10 cycles of the 40 ° errorclamp block. Each dot represents the data of one participant. The height of the bars represents the inter-individual means and the error bars represent the standard error of the mean.

We observed an age-related increase of internal model recalibration as measured with task-irrelevant feedback, which replicates our previous work (Vandevoorde and Orban de Xivry, 2019). In contrast to paradigm 1, performance for the position-matching task was reduced with aging for the two preregistered variables in paradigm 2 (Analysis 2; Figure 6A-B: VarXY: t(68) = - 3.6, p = 0.0006, d = - 0.87, CI95-d = [- 1.37, - 0.38]; AbsErXY: t(68) = - 2.5, p = 0.01, d = - 0.60, CI95-d = [ - 1.09, - 0.12]). However, there was no evidence that the slope or the inflection point differed across age groups (Analysis 3; slope: Figure 6C: Age: F(1,66) = 0.14, p = 0.71; η^2^ = 0.022; inflection point: Figure 6D: Age: F(1,66) = 2.60, p = 0.11; η^2^ = 0.037). Similar results were obtained for the interaction between age and the time point of the perceptual boundary test. No interaction effect between age and time point was observed for the slope of the psychometric curve (Analysis 3; Figure 6C: Age x Time: F(1,66) = 0.1, p = 0.75; η^2^ = 0.0059), and the inflection point (Analysis 3; Figure 6D: Age x Time: F(1,66) = 2.15, p = 0.15; η^2^ = 0.15).

**Figure 6:**
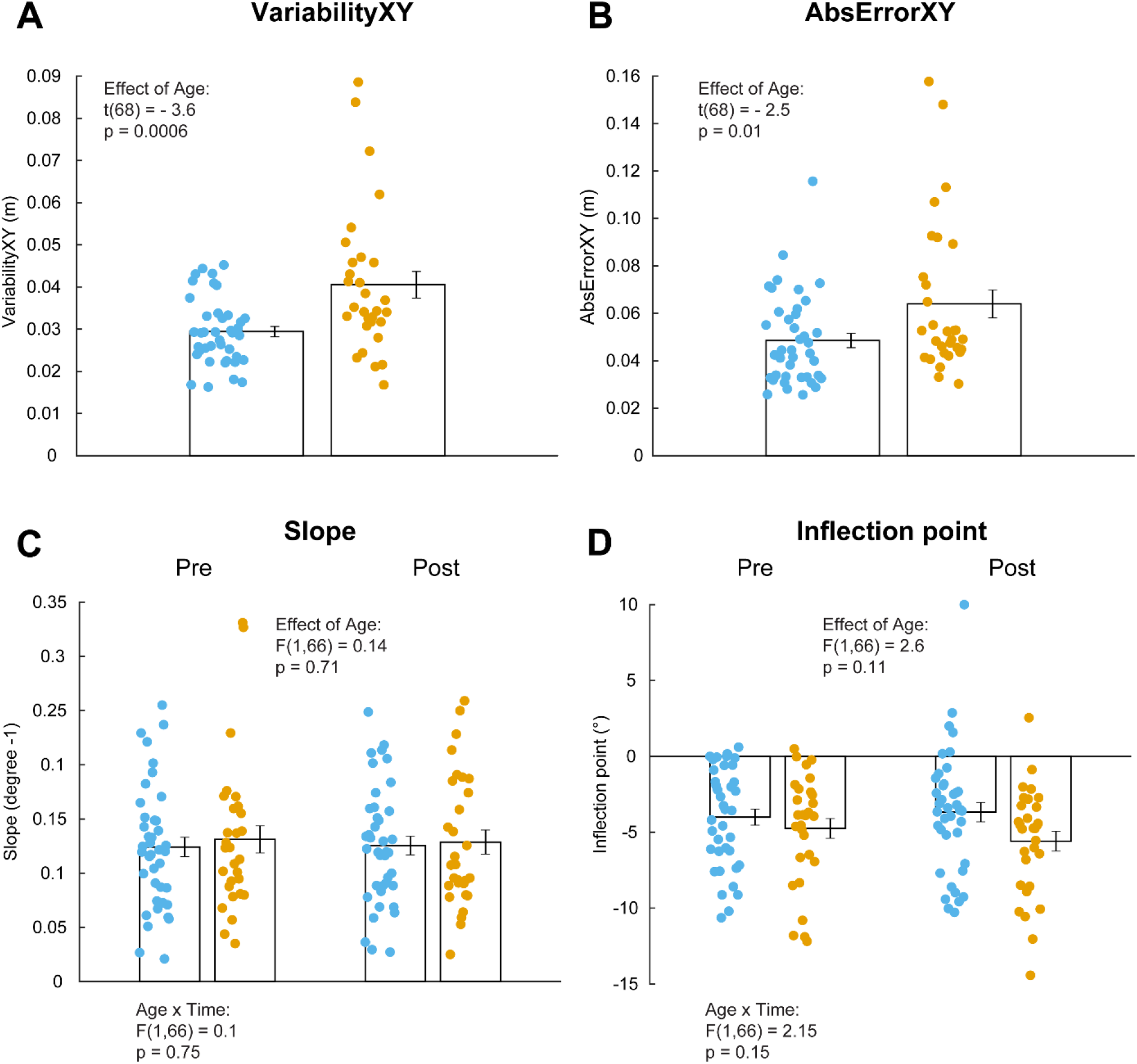
Age-related decline of somatosensation, visualized with data from participants of paradigm 2. **A-B:** Outcomes of the position matching task measured before adaptation: variability of hand position, VariabilityXY, and the absolute error, AbsErrXY. Lower values are associated with better somatosensory acuity. **C-D:** Outcomes of the perceptual boundary estimation task measured before (Pre) and after adaptation (post). For panel C, higher values are associated with better somatosensory acuity. No such interpretation exists for the inflection point. Each dot represents the data from one participant. The height of the bars represents the inter-individual means and the error bars represent the standard error of the mean.

Taking the direction of the perturbation into account, we did not find evidence for a difference in slope between pre and post (Analysis 3, F(1,68) = 0.00042, p = 0.98) or in inflection point (Analysis 3; F(1,68) = 0.39, p = 0.54). In addition, we failed to find evidence for a relationship between our level of internal model recalibration and change in inflection point between pre and post condition (Analysis 6; implicit adaptation: beta = 0.005; p = 0.5; Figure 8). Since we performed the perceptual boundary test after a short washout instead of before washout as in (Ostry et al., 2010), we verified whether the last cycles of the aftereffect were linked with the change in inflection point. However, no relationship existed between the aftereffect and change in inflection point between pre and post condition (Analysis 6; aftereffect: beta = 0.18; p = 0.1; Figure 8).

Next, the **preregistered** relationship between the position matching variables and implicit adaptation level from the task-irrelevant clamped feedback experiment was determined (Analysis 5). However, we did not find evidence that the position matching variables were related to the level of implicit adaptation (Figure 7A-B; Table 2; β_VAR-XY_ = - 0.08; p = 0.6, R = 0.12; β_ABS-XY_ = 0.04; p = 0.8, R = 0.02) or that perceptual boundary variables were related to the internal model recalibration. There was no evidence for a relationship between slope and implicit adaptation level (Figure 7C; β_SLOPE_ = - 0.15; p = 0.1, R = - 0.11). Yet, there was evidence for a negative link between inflection point and implicit adaptation (Figure 7D; pre: β_IP_ = −0.32, p = 0.001, R = - 0.37). However, it is difficult for us to relate somatosensory acuity and the inflection point. Furthermore, the inflection point was not different between young and old (Figure 6D), which makes it unclear how this variable could explain the age-related changes in implicit adaptation. Overall, no relation existed between somatosensatory acuity (measured by the position matching task or by the slope in the perceptual boundary task) and the level of implicit adaptation in both paradigms. Therefore, the **preregistered** hypothesis was not confirmed. Removing the two outliers in the older adult’s group did not change the outcome of our results. For completeness, the confidence intervals for all calculated regression coefficients are given (Figure 8).

**Figure 7:**
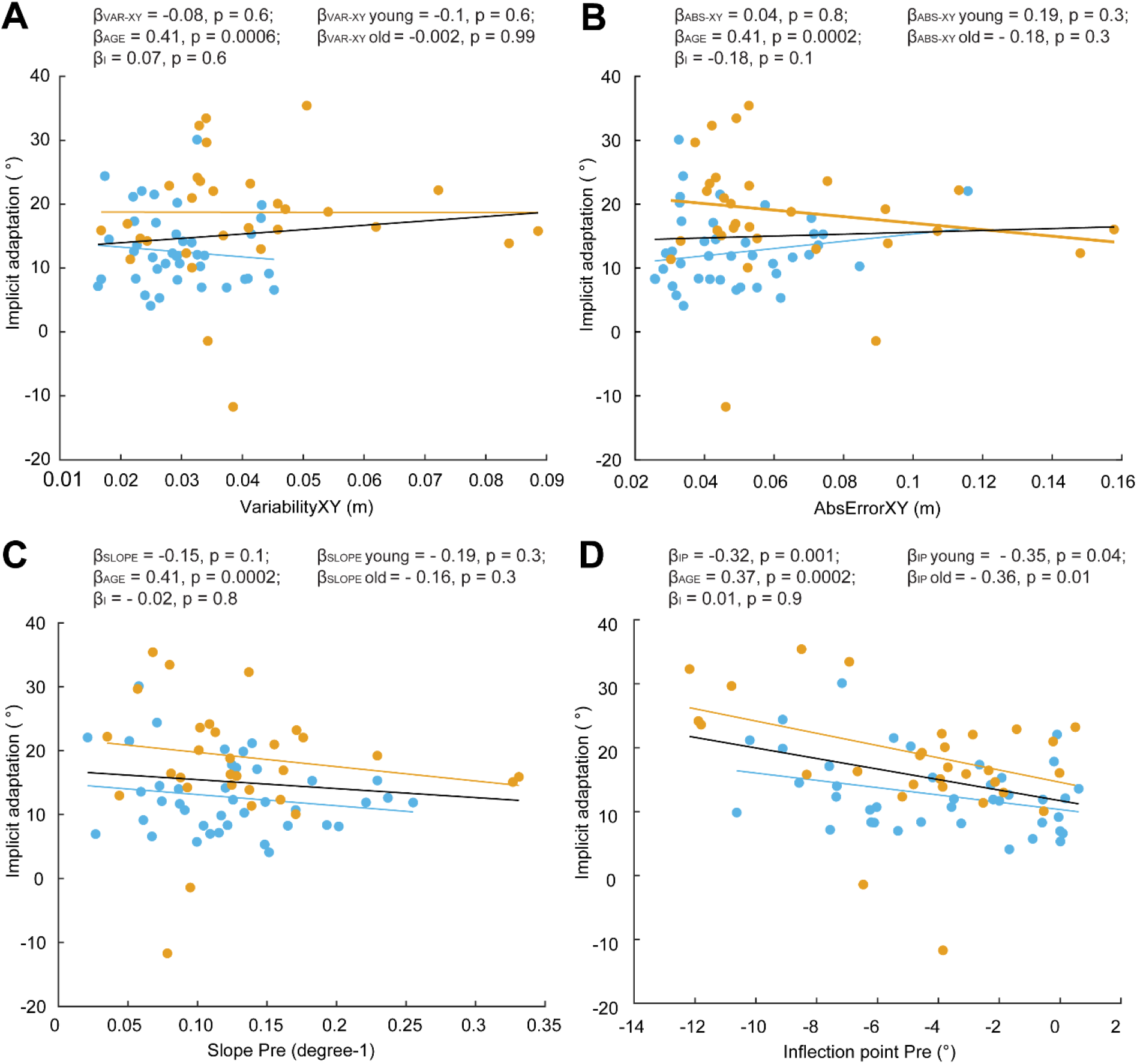
No relationship observed between implicit adaptation and somatosensory acuity. Data from paradigm 2. **A-B**: Scatterplot of the outcomes of the position matching task (data from fig.6A and B) and the extent of implicit motor adaptation from the task-irrelevant clamped feedback task (data from Fig.5B). A positive relationship is expected given that better somatosensory acuity is associated with lower values. **C-D**: Scatterplot of the outcomes of the perceptual boundary task (data from fig.6C and D) and the extent of implicit motor adaptation (data from Fig.5B). A negative relationship is expected for panel C given that better somatosensory acuity is associated with higher slope values. In all panels, each dot represent the data of a single participant. Solid lines represent the regression line from the robust regression analysis for both populations together (black line), for young participants only (blue lines) and for old participants only (orange lines).

Besides verifying the preregistered hypothesis, we could compare the different methods that we used to measure somatosensory acuity. The position matching task measures position sense, while the perceptual boundary test was a custom developed task measuring kinesthetic sense. Therefore, it is not known how these tasks would compare to each other. Data were obtained for both tasks in both paradigms, which allowed us to combine data of all 133 participants for this comparison. We compared the two preregistered variables of the position matching task to the two preregistered variables from the perceptual boundary test (Analysis 7; Figure 8). We observed a negative relationship between the slope variable of the perceptual boundary curve and VariabilityXY of the position matching task (Figure 8; β = −0.25; p = 0.009).

**Figure 8:**
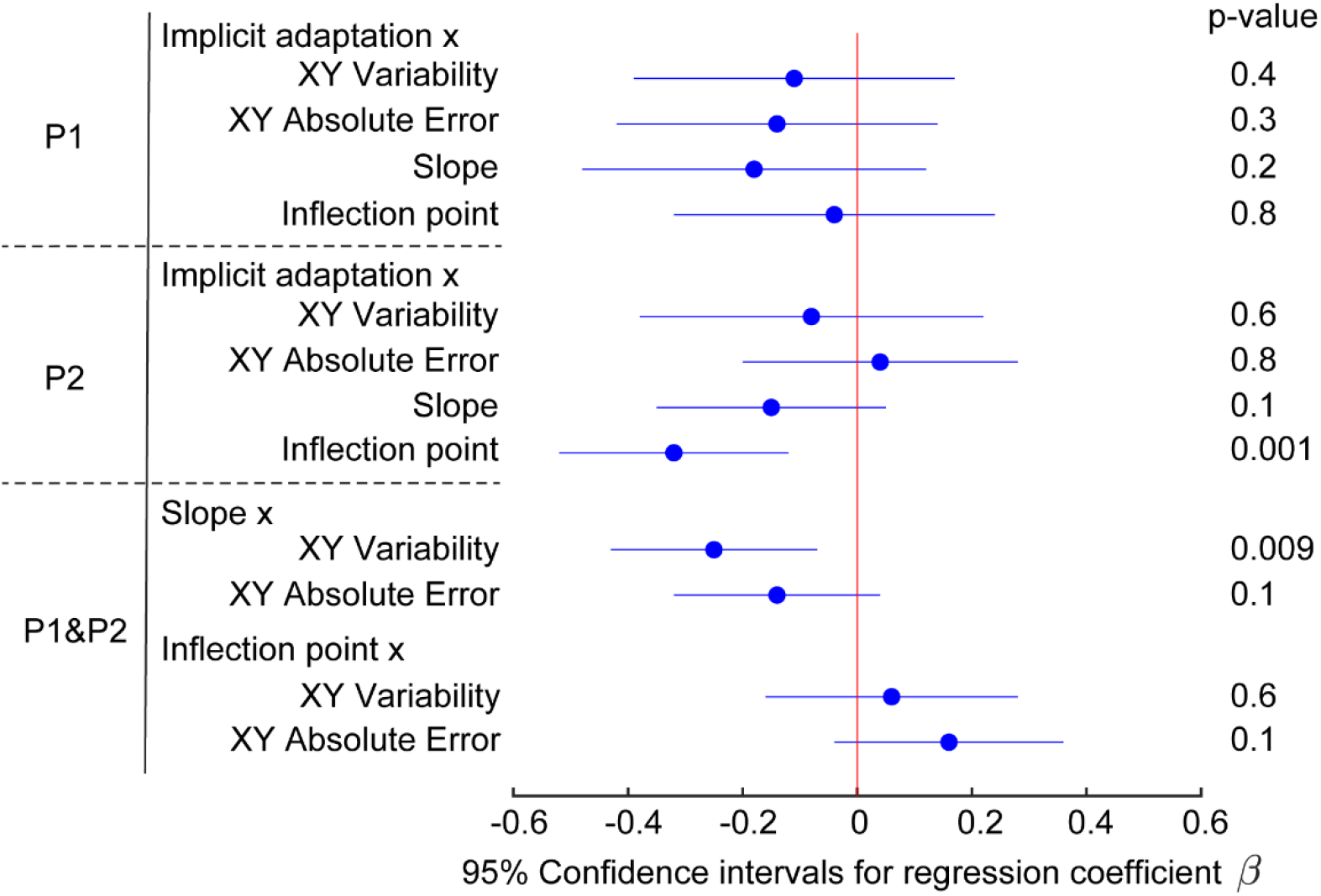
95% Confidence intervals for regression coefficient (β) from all analyses. The dots represent the mean and the error bar the confidence interval as assessed by bootstrapping. In the first two rows, the dependent variable is the extent of implicit adaptation and each row corresponds to different independent variables. Each row is associated with a different paradigm. In the third row, data from all participants are pooled together. Outcomes from the perceptual boundary task are used as dependent variables and those from the position-matching task are used as independent variables in the regression. In all regressions, age group is added as an additional independent factor. Numbers on the right correspond to the p-values associated with the different regression coefficients.

## Discussion

In this paper, we aimed to determine whether older adults’ increased recalibration of the internal model could be caused by a reduction of somatosensory acuity with aging. To that end, we quantified implicit adaptation with cued motor adaptation in a first paradigm, in which color cues indicated the presence of the perturbation (Morehead et al., 2015). However, we did not find any evidence that the level of implicit adaptation was related to somatosensory acuity in that experimental context. In a second paradigm, we assessed internal model recalibration independently of explicit strategy by providing cursor-irrelevant feedback (Morehead et al., 2017) and observed an increased internal model recalibration with aging which confirms our previous work (Vandevoorde and Orban de Xivry, 2019). According to our **preregistered hypothesis**, we expected a negative relationship between internal model recalibration and somatosensory acuity. However, we failed to support this hypothesis since there was no evidence for a relation between somatosensory acuity and internal model recalibration, measured in the two different paradigms.

Furthermore, we assessed somatosensory acuity as position and kinesthetic sense. An age-related decline of position sense could be observed in this second paradigm (Figure 6A-B). This difference was not detected in the first paradigm (Figure 3A-B), but can simply arise because of natural sampling variability. A difference in position sense replicates the observations that a decline in position sense accompanies healthy aging (Herter et al., 2014). On the contrary, an age-related decline of kinesthetic sense was not observed (Figure 3C-D, Figure 6C-D).

### Somatosensation and internal model recalibration

Studies of deafferented patients show that proprioception is important to maintain an accurate internal model as these patients exhibit impaired inter-segmental coordination. Such coordination relies presumably on internal model function in healthy participants (Ghez et al., 1990; Sainburg et al., 1995). It is therefore hypothesized that arm proprioception might contribute to the maintenance of an accurate internal model (Scarchilli et al., 1999). While a previous study had reported no influence of somatosensory impairment on adaptation to a visuomotor rotation (Bernier et al., 2006), that paper did not assess pure implicit adaptation and these results could have been confounded by the presence of a compensatory explicit strategy. This is why we believe that, if somatosensation was important to maintain the accuracy of the internal model, age-related somatosensory deficits could account for the paradoxical increase in internal model recalibration observed in our previous study (Vandevoorde and Orban de Xivry, 2019).

Yet, we failed to observe any consistent evidence in favor of that hypothesis. In paradigm 1, there was a slightly larger implicit adaptation for elderly participants compared to younger ones but we did not find any difference in somatosensory function. In paradigm 2, we found some differences in somatosensation between the two age groups (position sense was reduced in older compared to younger adults), while the difference in implicit adaptation was quite large with the elderly exhibiting much more implicit adaptation than young participants. Finally, we failed to find a relationship between somatosensory function and implicit adaptation as could have been expected from studies on deafferented patients (Ghez et al., 1990; Sainburg et al., 1995), which showed that somatosensory function is important for internal model recalibration.

While not significant, the β-coefficient for regression between somatosensory acuity (slope parameter) and the extent of implicit motor adaptation was around - 0.2 in both experiments. There is thus the possibility that a significant relationship could be found with many more participants. Yet, it is difficult to conceive that such a weak relationship between somatosensory function and implicit adaptation could explain the large age-related differences in implicit adaptation found in paradigm 2 and in our previous paper (Vandevoorde and Orban de Xivry, 2019).

Recalibration of proprioception goes along with recalibration of the internal model. Several papers have demonstrated that visuomotor discrepancies (the difference between the visual feedback provided and the direction of hand motion) yield both changes in hand localization and in internal model recalibration and that these two quantities are linked to each other (Gastrock et al., 2020; Salomonczyk et al., 2013; Tsay et al., 2020). Interestingly, this also holds in older people where proprioceptive recalibration and after-effects are comparable to younger people (Cressman et al., 2010).

Yet, one could expect that inter-individual differences in proprioceptive acuity could account for differences in proprioceptive or internal model recalibration. Indeed, if visual and proprioceptive information are combined in an optimal manner (van Beers et al., 2002, 1999), a visuo-proprioceptive discrepancy (a difference between the direction of the hand and the direction of the cursor on the screen) is typically resolved by recalibrating the less reliable source of information, here proprioception. If proprioception is less reliable such as in older people (Goble et al., 2009), one could expect that it would be recalibrated even further for that population. Yet, recalibration of proprioception is comparable in young and old participants (Cressman et al., 2010) despite the fact that uncertainty in hand localization was larger in older people.

In visuomotor perturbation, internal model recalibration in older people is also comparable to or larger than that of younger people (Cressman et al., 2010; Hegele and Heuer, 2013; Heuer and Hegele, 2008; Reuter et al., 2020, 2018; Vachon et al., 2020; Vandevoorde and Orban de Xivry, 2019). We were expecting that the visuo-proprioceptive discrepancy would be responsible for the amplitude of the adaptative response. Over a single trial, the amplitude of the adaptative response grows linearly with the size of the observed error over a small range and then saturates for larger errors (Hutter and Taylor, 2018; Kim et al., 2018; Marko et al., 2012). It was suggested that the saturation of the adaptation response could be due to the discrepancy between visual and proprioceptive estimates (Wei and Kording, 2009) where large discrepancies are considered irrelevant for the motor system. The evaluation of this discrepancy requires reliable visual and proprioceptive signals. Unreliable proprioceptive information (e.g. due to aging) should increase the size of errors deemed as relevant and to which the motor system needs to respond and should also yield to a larger adaptative response. Following this theory, the quality of proprioception should be negatively linked to the size of the discrepancies that are deemed relevant and to which the motor system respond. The present experiments did not provide any evidence for this negative link. A similar result has been recently observed with a force-field perturbation (Kitchen and Miall, 2020) where it was found that dynamic proprioceptive acuity, which was measured similarly as we did, was not related to the extent of force-field after-effect in young and old people. Together, these results and ours cast doubts on the fact that vision and proprioception are combined to drive motor adaptation or other age-related effects on motor function such as larger sensory attenuation (Parthasharathy et al., 2020).

This finding can be put into the larger debate about the possible combination of vision and proprioception during reaching or motor adaptation tasks. Optimal combination has been hypothesized on the basis of different amount of adaptation across direction (van Beers et al., 2002). In contrast, several papers demonstrate that the two sensory signals contribute differently to sensory motor adaptation. For instance, Hayashi and colleagues (2020) demonstrated that visual and proprioceptive feedback were linked to separate motor memories. Furthermore, Scheidt et al. (2005) showed that adaptation with and without vision was identical but that adaptation to the force-field perturbation was absent when the visual feedback of the cursor was constrained to a straight line going right to the target. This suggests that, during adaptation, visual feedback (the hand going straight) dominated the proprioceptive information about the perturbation (see also Judkins and Scheidt, 2014). Similarly, disrupting proprioception via muscle vibration does not have any effect on the adaptation to a visuomotor rotation where there is a discrepancy between vision and proprioception because, again, vision dominates the adaptation response (Pipereit et al., 2006). Our results are in line with the fact that, during motor adaptation tasks, vision and proprioception are not optimally combined. Note that such optimal integration remains valid during perceptual tasks (Ernst and Banks, 2002; van Beers et al., 1999).

### Alternative explanations for increased internal model recalibration with aging

We failed to find evidence that an age-related decline in somatosensation with aging was related to internal model recalibration increase with aging. This suggests that internal model recalibration increases with aging, independent from somatosensory deficits. We initially suggested two explanations for the increased internal model recalibration related to somatosensory acuity changes: 1) an altered sensory integration with an up weighting of visual feedback because of reduced somatosensory acuity, and 2) an increased reliance on predicted sensory outcome because of reduced somatosensory acuity. However, these two initial explanations seem unlikely given our results and alternative explanations should be explored.

First, it could be that visual acuity is the relevant sensory information source for internal model recalibration of arm reaching, and not somatosensory acuity. For that reason, it remains interesting to investigate the relationship between visual acuity and the level of internal model recalibration for a group of young and older adults. This explanation might be related to studies investigating sensory attenuation. Sensory attenuation is the decreased sensation of a self-produced stimulus (Blakemore et al., 2000; Wolpert et al., 1995). Two studies (Parthasharathy et al., 2020; Wolpe et al., 2016) observed an age-related increase of sensory attenuation for force applied to the finger and the upper-limb. This shift in sensory attenuation with aging resulted in increased overcompensation to match a previously applied force. Wolpe et al. (2016) suggested that this shift in sensory attenuation was related to a reduced somatosensory acuity with aging but this was not confirmed by later experiments studies (Parthasharathy et al., 2020). It is not directly related to our observation of increased internal model recalibration with aging, since we applied visual instead of force perturbations. Nevertheless, it is important to verify whether sensory attenuation is not involved in our observation as well. For instance, if the stronger reaction to visual error with aging is related to a reduced visual acuity.

A second explanation might be an increased internal model recalibration because of life-long learning from sensory prediction errors. Throughout life, errors of all kind of sizes are experienced, likely creating a growing memory of sensory prediction errors and appropriate responses to them, which could upregulate error-sensitivity with aging (Albert et al., 2020; Herzfeld et al., 2014). This increased internal model recalibration might have arisen as a compensation mechanism for a reduction of explicit aiming with aging (Heuer and Hegele, 2008; Vandevoorde and Orban de Xivry, 2020, 2019); this compensation mechanism has also been observed outside an aging context (Bond and Taylor, 2015; Brudner et al., 2016; Christou et al., 2016; Schween and Hegele, 2017; Taylor and Ivry, 2011). Our experiment specifically investigated angular sensory prediction errors. However, it would be useful to verify whether elderly react more to other types of sensory prediction errors such as errors induced by small random force-field perturbations where no explicit aiming strategies can be applied. Future studies should try to determine whether elderly react more to other sensory prediction errors as well, such as observed in gait, speech or force-field adaptation.

A third explanation is that older adults are less able to filter the distracting irrelevant visual feedback. Elderly are exhibiting impaired selective attention, which results in lower abilities to enhance task-relevant information and suppress task-irrelevant information (Awh et al., 2006; Gazzaley and Nobre, 2012). Applied to increased internal model recalibration, this might result in lower abilities to suppress the task-irrelevant visual feedback and enhance the relevant somatosensory feedback. According to this explanation, no link is required between increased internal model recalibration and (somato)sensory acuity but instead a link might exist between internal model recalibration and working memory assessment that incorporates the ability to filter irrelevant stimuli (McNab and Klingberg, 2008).

A fourth possible explanation is that the reward structure might have a different effect on young and older adults. The absence of reward is known to drive the implicit system, while the presence of reward is reducing it (Kim et al., 2019). In our experiments, the absence of reward could have resulted in a bigger driving force for the implicit system of older adults compared to younger adults.

Finally, it is possible that conscious reports of somatosensation are irrelevant for motor adaptation because different proprioceptive signals are used from motor control and for perception. For instance, Rand and Heuer (2020) recently that conscious reports of visual and proprioceptive information was not consistent with sensory integration while unconscious proprioceptive signals reflected in the direction of reaching movements were clearly integrated with visual signals. In this case, conscious reports of hand position that are influenced by experimental manipulations like visuomotor rotation (Cressman and Henriques, 2011), might be completely independent of the proprioceptive signals used by the motor system. Such dissociation between conscious perception and action has been reported before for the size-weight illusion (Flanagan and Beltzner, 2000).

### Age-related deficits in somatosensation are limited

The position matching task is a reliable, quantitative tool for multijoint upper limb position sense (Dukelow et al., 2010) with an excellent reliability and interrater variability for multijoint limb position sense. The second somatosensory test consisted of a robotic assessment of kinesthetic sense, which had been used before with horizontal (Kitchen and Miall, 2018; Ostry et al., 2010) and angular displacements (Darainy et al., 2013). This test is sensitive enough to detect changes in proprioception induced by force-field adaptation (Ostry et al., 2010). Our measurements were limited to active limb movement since this mimicked best the motor learning paradigm requirements.

One limitation of the kinesthetic somatosensory test was the requirement of an overt response since somatosensation is rather a covert process. However, the possibility to guess in case of doubts revealed that the participants were quite accurate despite being clueless. For instance, the examiner (KV) noticed that, although participants answered correctly to the deviation, they often responded that they had no clue whether their answer was correct. An improvement to the task could be to allow participants to give two more answer choices (‘maybe left’, ‘maybe right’) or to report the certainty of their response on a scale. The outcomes of both tests could be influenced by central processing abilities and working memory, which are typically reduced with aging. However, we instructed participants clearly and allowed them to practice long enough until they understood the tasks correctly. Therefore, we think that the influence of cognitive decline is negligible in our assessment.

We expected that somatosensory acuity would decline with aging in both sensory tests. This expectation appears partially correct for position sense only, while it was not for kinesthetic sense since the slope of the psychometric accuracy curve was similar for young and older adults. Previous literature about kinesthetic sense and aging is limited and inconclusive. One study (Kitchen et al., 2019) reports an age-dependent increase in proprioceptive bias in active multi-joint upper limb movement but not in uncertainty range. In contrast, another study (Cressman et al., 2010) reports a similar bias between age groups but a slightly increased uncertainty range in elderly. In future experiments, it would be interesting to clarify these differences. One might design a more sensitive kinesthetic paradigm, for instance by applying a fixed time duration for each reaching trial, by controlling grip strength or by increasing the number of trials and angular deviations. Finally, one could replace the PEST algorithm by the likely more sensitive Bayesian adaptative QUEST algorithm (Watson and Pelli, 1983). However, if an effect between groups exists, it will be small and it is probably not related to age-related deficits in motor adaptation.

## Conclusion

Based on two different motor adaptation experiments, we were able to confirm that internal model recalibration is not impaired with aging but is increased when isolated from the explicit process. However, we failed to support the hypothesis that a decline in somatosensory acuity could explain this increase in internal model recalibration with age. This questions the fact that vision and somatosensation are dynamically combined during motor adaptation. There is a need to incorporate these age-related effects in the current models of sensorimotor adaptation in order to transition from fundamental insights in motor learning to clinical applications.

## Acknowledgements

This work was supported by an internal grant of the KU Leuven (STG/14/054) and by the FWO (1519916N).

